# Multiple Pathways Impact Swarming Motility of *Pseudomonas fluorescens* Pf0-1

**DOI:** 10.1101/2024.01.17.576057

**Authors:** Alexander B. Pastora, Kara M. Rzasa, George A. O’Toole

## Abstract

Swarming motility in pseudomonads typically requires both a functional flagellum and production/secretion of a biosurfactant. Published work has shown that the wild-type *Pseudomonas fluorescens* Pf0-1 is swarming-deficient due to a point mutation in the *gacA* gene, which until recently, was thought to inactivate rather than attenuate the Gac/Rsm pathway. As a result, little is known about the underlying mechanisms that regulate swarming motility by *P. fluorescens* Pf0-1. Here, we demonstrate that a Δ*rsmA* Δ*rsmE* Δ*rsmI* mutant, which phenotypically mimics Gac/Rsm pathway overstimulation, is proficient at swarming motility. RsmA and RsmE appear to play a key role in this regulation. Transposon mutagenesis of the Δ*rsmA* Δ*rsmE* Δ*rsmI* mutant identified multiple factors that impact swarming motility, including pathways involved in flagellar synthesis and biosurfactant production/secretion. We find that loss of genes linked to biosurfactant Gacamide A biosynthesis or secretion impact swarming motility, as does loss of the alternative sigma factor FliA, which results in a defect in flagellar function. Collectively, these findings provide evidence that *P. fluorescens* Pf0-1 can swarm if the Gac/Rsm pathway is activated, highlight the regulatory complexity of swarming motility in this strain, and demonstrate that the cyclic lipopeptide Gacamide A is utilized as a biosurfactant for swarming motility.

**Importance:** Swarming motility is a coordinated process that allows communities of bacteria to collectively move across a surface. For *P. fluorescens* Pf0-1, this phenotype is notably absent in the parental strain and to date little is known about the regulation of swarming in this strain. Here, we identify RsmA and RsmE as key repressors of swarming motility via modulating the levels of biosurfactant production/secretion. Via transposon mutagenesis and subsequent genetic analyses, we further identify potential regulatory mechanisms of swarming motility and link Gacamide A biosynthesis and transport machinery to swarming motility.

## Introduction

Swarming motility is a highly coordinated form of movement, generally requiring a functional flagellum and a wetting agent, that allows for the rapid translocation of surface-associated bacterial communities. Swarming has been characterized in a variety of gram-positive and gram-negative bacteria including *Bacillus subtilis*, *Escherichia coli*, *Pseudomonas aeruginosa*, *Salmonella enterica*, and *Serratia marcescens* (1–15). Regulation of swarming motility has been extensively studied in the pseudomonads (16–28).

For *Pseudomonas fluorescens*, swarming motility is important for efficient colonization of plant roots and rapid movement towards root exudates (29–32). Interestingly, the wild-type *P. fluorescens* Pf0-1 strain, which was isolated from nutrient rich sandy-loam soil (33), is deficient for swarming motility due to a N109P point mutation in the *gacA* gene, which was previously thought to completely abolish function of the Gac/Rsm pathway (34, 35). However, a recent study determined that the *Pseudomonas fluorescens* Pf0-1 Gac/Rsm pathway could be stimulated by overproduction of the upstream histidine kinase GacS, suggesting that the N109P mutation in *gacA* gene attenuates rather than abolishes the Gac/Rsm pathway of *P. fluorescens* Pf0-1, and that overstimulation of the pathway could be achieved via chromosomal deletions of *rsmA*, *rsmE*, and *rsmI* genes (36).

Here we show that the Δ*rsmA* Δ*rsmE* Δ*rsmI* triple mutant of *P. fluorescens* Pf0-1 is capable of swarming motility. We demonstrate that loss of RsmA and RsmE functions is sufficient to induce swarming by *P. fluorescens* Pf0-1. Starting with the Δ*rsmA* Δ*rsmE* Δ*rsmI* mutant, we use a genetic screen and subsequent molecular genetic studies to identify pathways involved in swarming motility. These pathways impact flagellar motility and link the biosynthesis of the biosurfactant Gacamide A and its secretion machinery with swarming motility.

## Results

### The Gac/Rsm pathway regulates swarming motility in an RsmA- and RsmE-dependent manner

To assess the role of the Gac/Rsm pathway for swarming motility, we utilized a strain deficient in the small proteins RsmA, RsmE, and RsmI (Rsm proteins) in *P. fluorescens* Pf0-1, which was previously demonstrated to mimic overstimulation of the Gac/Rsm Pathway (36). We probed this strain for flagellar function using a Tris-buffered minimal medium supplemented with L-arginine as the sole carbon source (KA medium) supplemented with 0.3% agar (designated “swim agar” for assessing swimming motility) or 0.5% agar (designated “swarm agar” for assessing swarming motility).

As previously shown, low percentage agar media, including swarm agar, can also be used to probe for biosurfactant production/secretion by the pseudomonads. Biosurfactant production is phenotypically defined as a translucent liquid zone radiating from the inoculum that then forms the leading edge of the subsequent swarm zone (11, 37, 38). In this study, we designate this phenotype on swarm agar as the “biosurfactant zone”. As a negative control, we included a strain deficient in FleQ function (a 11*fleQ* mutant), which has previously been shown to lack flagellar motility in multiple pseudomonads (24, 27, 39–42).

After 24h growth on swim agar at 30°C, the Δ*rsmA* Δ*rsmE* Δ*rsmI* triple mutant, which was capable of swimming, shows significantly less swimming motility compared to the wild-type (WT) strain (Figure 1A). This reduced swimming phenotype was reported previously by Pastora and O’Toole (36) and attributed to differences in planktonic growth (36). In contrast, the Δ*fleQ* and Δ*fleQ* Δ*rsmA* Δ*rsmE* Δ*rsmI* mutant strains were completely deficient for swimming motility (Figure 1A).

**FIG 1.**
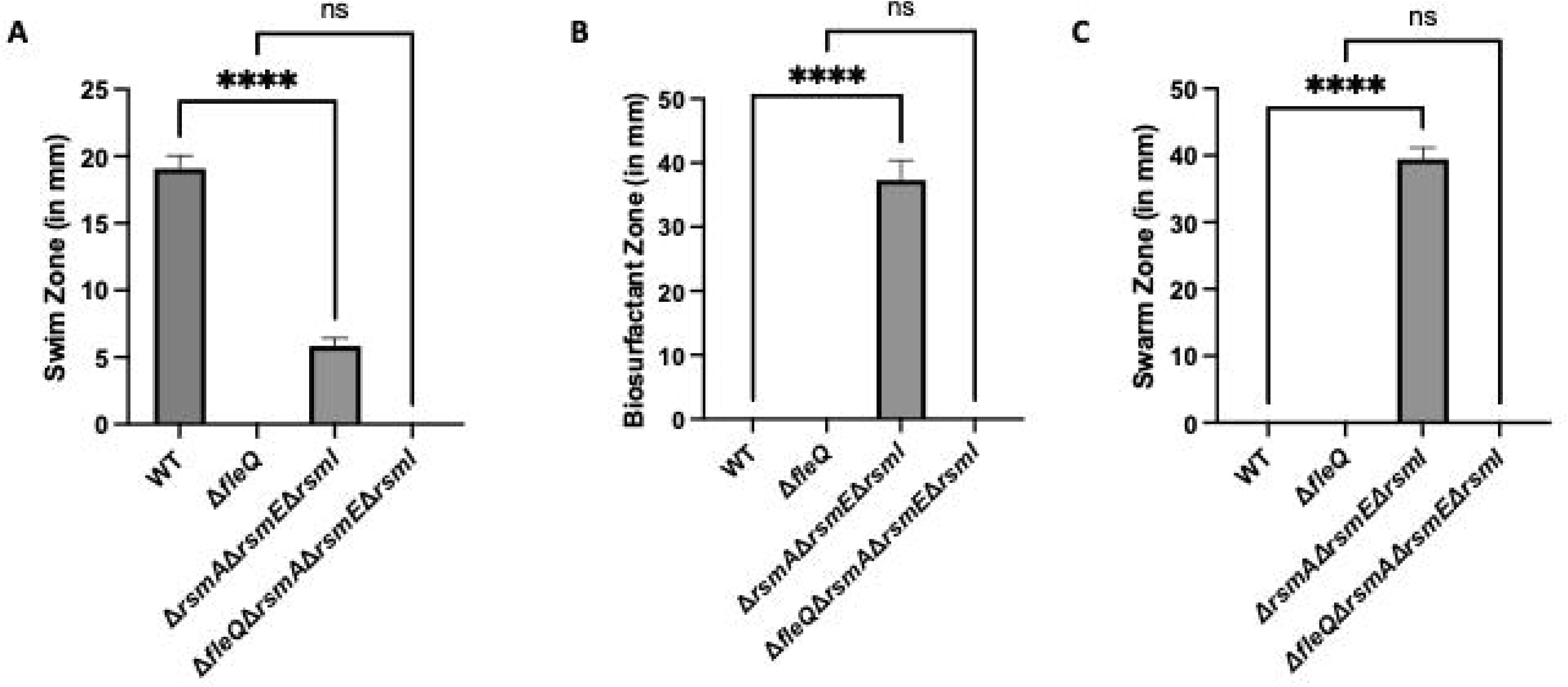
A strain deficient for the Rsm proteins swarms. (A) Swim zone (in millimeters) of the WT strain, Δ*fleQ* single mutant, Δ*rsmA* Δ*rsmE* Δ*rsmI* triple mutant, and a Δ*fleQ* Δ*rsmA* Δ*rsmE* Δ*rsmI* quadruple mutant after toothpick inoculation on KA minimal medium supplemented with 0.3% agar followed by 24h growth at 30°C. (B) Biosurfactant zone (in millimeters) of the WT strain, Δ*fleQ* single mutant, Δ*rsmA* Δ*rsmE* Δ*rsmI* triple mutant, and a Δ*fleQ* Δ*rsmA* Δ*rsmE* Δ*rsmI* quadruple mutant after inoculation of 2.5µl of overnight culture on the surface of KA minimal medium supplemented with 0.5% agar followed by 24h growth at 30°C. (C) Swarm zone (in millimeters) of the WT strain, Δ*fleQ* single mutant, Δ*rsmA* Δ*rsmE* Δ*rsmI* triple mutant, and a Δ*fleQ* Δ*rsmA* Δ*rsmE* Δ*rsmI* quadruple mutant after inoculation of 2.5µl of overnight culture on the surface of KA minimal medium supplemented with 0.5% agar followed by 24h growth at 30°C then 24h growth at room temperature. Statistical significance for this figure was determined using one-way ANOVA with Tukey’s multiple comparisons tests. ****, P<0.0001. All error bars represent standard deviation.

After 24h growth on swarm agar at 30°C, the Δ*rsmA* Δ*rsmE* Δ*rsmI* mutant produced a biosurfactant zone while the WT strain and FleQ-deficient mutants did not produce a measurable biosurfactant zone (Figure 1B). Since this incubation time was not sufficient to induce swarming motility in any of the tested strains, we subsequently incubated the strains for an additional 24h at room temperature. After 24h growth on swarm agar at 30°C then 24h growth at room temperature, the Δ*rsmA* Δ*rsmE* Δ*rsmI* mutant displayed motility on swarm agar, whereas the other strains were deficient for swarming motility (Figure 1C). Interestingly, the Δ*rsmA* Δ*rsmE* Δ*rsmI* mutant both swims, produces biosurfactant, and eventually swarms on swim agar (Figure S1), although to a lesser extent than on swarm agar.

To identify the Rsm proteins associated with swarming motility, we made chromosomal deletions of the individual genes coding for the Rsm proteins in the WT and FleQ-deficient background strains and assessed these strains on swarm agar. Loss of any individual Rsm protein was not sufficient to induce swarming motility (Figure 2A). Analysis of these strains for swimming motility and biosurfactant production revealed that while the single *rsm* deletions in the WT background were proficient for swimming motility (Figure S2A), none of these strains were able to produce a biosurfactant zone (Figure S2B).

**FIG 2.**
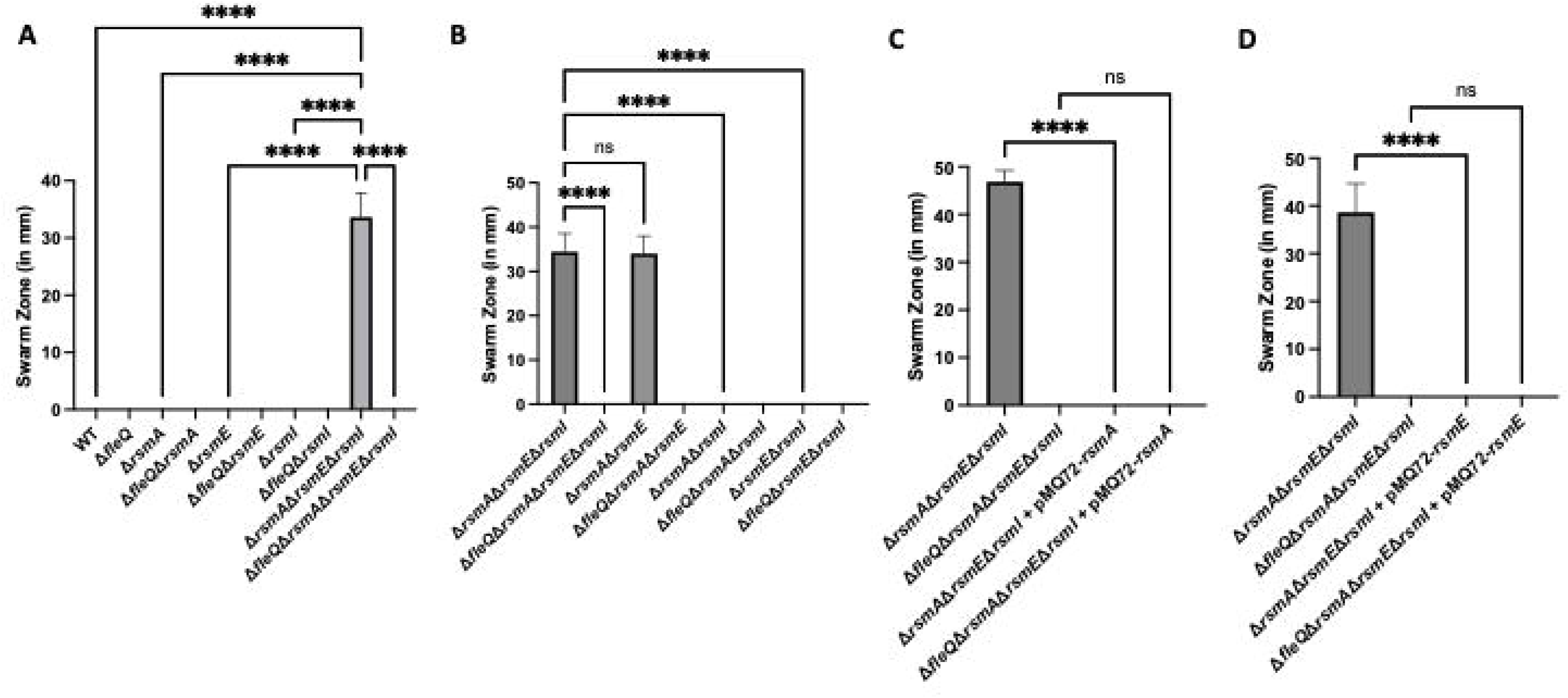
A strain deficient for both RsmA and RsmE shows swarming motility. (A) Swarm zone (in millimeters) of the WT strain, Δ*fleQ* single mutant, Δ*rsmA* single mutant, Δ*fleQ* Δ*rsmA* double mutant, Δ*rsmE* single mutant, Δ*fleQ* Δ*rsmE* double mutant, Δ*rsmI* single mutant, Δ*fleQ* Δ*rsmI* double mutant, Δ*rsmA* Δ*rsmE* Δ*rsmI* triple mutant, and a Δ*fleQ* Δ*rsmA* Δ*rsmE* Δ*rsmI* quadruple mutant after inoculation of 2.5µl of overnight culture on the surface of KA minimal medium supplemented with 0.5% agar followed by 24h growth at 30°C then 24h growth at room temperature. (B) Swarm zone (in millimeters) of the Δ*rsmA* Δ*rsmE* Δ*rsmI* triple mutant, Δ*fleQ* Δ*rsmA* Δ*rsmE* Δ*rsmI* quadruple mutant, Δ*rsmA* Δ*rsmE* double mutant, Δ*fleQ* Δ*rsmA* Δ*rsmE* triple mutant, Δ*rsmA* Δ*rsmI* double mutant, Δ*fleQ* Δ*rsmA* Δ*rsmI* triple mutant, Δ*rsmE* Δ*rsmI* double mutant, and Δ*fleQ* Δ*rsmE* Δ*rsmI* triple mutant after inoculation of 2.5µl of overnight culture on the surface of KA minimal medium supplemented with 0.5% agar followed by 24h growth at 30°C then 24h growth at room temperature. (C) Swarm zone (in millimeters) of the Δ*rsmA* Δ*rsmE* Δ*rsmI* mutant, Δ*fleQ* Δ*rsmA* Δ*rsmE* Δ*rsmI* mutant, Δ*rsmA* Δ*rsmE* Δ*rsmI* mutant + pMQ72-*rsmA*, and Δ*fleQ* Δ*rsmA* Δ*rsmE* Δ*rsmI* mutant + pMQ72-*rsmA* after inoculation of 2.5µl of overnight culture on the surface of KA minimal medium supplemented with 0.5% agar followed by 24h growth at 30°C then 24h growth at room temperature. (D) Swarm zone (in millimeters) of the Δ*rsmA* Δ*rsmE* Δ*rsmI* mutant, Δ*fleQ* Δ*rsmA* Δ*rsmE* Δ*rsmI* mutant, Δ*rsmA* Δ*rsmE* Δ*rsmI* mutant + pMQ72-*rsmE*, and Δ*fleQ* Δ*rsmA* Δ*rsmE* Δ*rsmI* mutant + pMQ72-*rsmE* after inoculation of 2.5µl of overnight culture on the surface of KA minimal medium supplemented with 0.5% agar followed by 24h growth at 30°C then 24h growth at room temperature. Statistical significance for this figure was determined using one-way ANOVA with Tukey’s multiple comparisons tests. ****, P<0.0001. All error bars represent standard deviation. All error bars represent standard deviation.

We made combinatorial chromosomal deletions of the genes coding for the various Rsm proteins in the WT and FleQ-deficient backgrounds and assessed these mutants for their ability to swarm. After growth for 24h at 30°C then 24h at room temperature, the Δ*rsmA* Δ*rsmE* double mutant was able to swarm to levels similar to the Δ*rsmA* Δ*rsmE* Δ*rsmI* triple mutant, while the other combinatorial deletions were deficient for swarming motility (Figure 2B). Analysis of these strains for swimming motility and biosurfactant production revealed that all deletions in the WT background were proficient for swimming motility (Figure S3A), but of these strains, only the Δ*rsmA* Δ*rsmE* double and Δ*rsmA* Δ*rsmE* Δ*rsmI* triple mutants were proficient for biosurfactant production (Figure S3B).

To further demonstrate the role of RsmA and RsmE in regulating swarming motility, we inserted the coding sequence of RsmA or RsmE into the arabinose-inducible shuttle vector pMQ72 directly downstream of the P*_BAD_* promoter (see Materials and Methods), transformed the constructs into the Δ*rsmA* Δ*rsmE* Δ*rsmI* and Δ*fleQ*Δ*rsmA* Δ*rsmE* Δ*rsmI* backgrounds and assessed these strains for their ability to swarm. After growth for 24h at 30°C then 24h at room temperature on swarm agar without inducing conditions (no addition of arabinose), the Δ*rsmA* Δ*rsmE* Δ*rsmI* + pMQ72-*rsmA* and Δ*rsmA* Δ*rsmE* Δ*rsmI* + pMQ72-*rsmE* strains are completely deficient for swarming motility (Figure 2C and 2D, respectively). Analysis of these strains revealed that Δ*rsmA* Δ*rsmE* Δ*rsmI* + pMQ72-*rsmA* and Δ*rsmA* Δ*rsmE* Δ*rsmI* + pMQ72-*rsmE* strains were proficient in swimming motility (Figures S4A and S4B), and these strains had a statistically significant decrease in biosurfactant secretion compared to their parental strains (Figure S4C and S4D).

### Transposon mutagenesis reveals genes required for swarming motility

Given our observation that the Δ*rsmA* Δ*rsmE* Δ*rsmI* strain is proficient for swarming motility, we generated a Tn*M* mariner transposon mutant library as previously described in Pastora and O’Toole (36). We initially probed this library for candidates that were able to swarm after 24h of growth at 30°C on swarm agar (hyper-swarm candidates) or were completely deficient in swarming motility after 24h of growth at 30°C then 24h of growth at room temperature on swarm agar (swarm-deficient candidates) and identified the approximate transposon insertion site and P*_TAC_* promoter orientation using arbitrary primed PCR.

These candidates were re-screened for swimming motility, biosurfactant production, and swarming motility. All candidates are listed in Table S1 with the associated locus tag, gene name, GenBank description, KEGG BRITE Terms, and KEGG Pathway Terms (where applicable). The candidates were also grouped separately into hyper-swarm candidates (Table S2) and swarm-deficient candidates (Table S3). The swarm-deficient candidates were additionally grouped independently based on loss of biosurfactant production (Table S4) or loss of swimming motility (Table S5; note some of the candidates are listed in more than one table). The candidates in each table were categorized based on annotated function and are graphically summarized in Figure 3.

**FIG 3.**
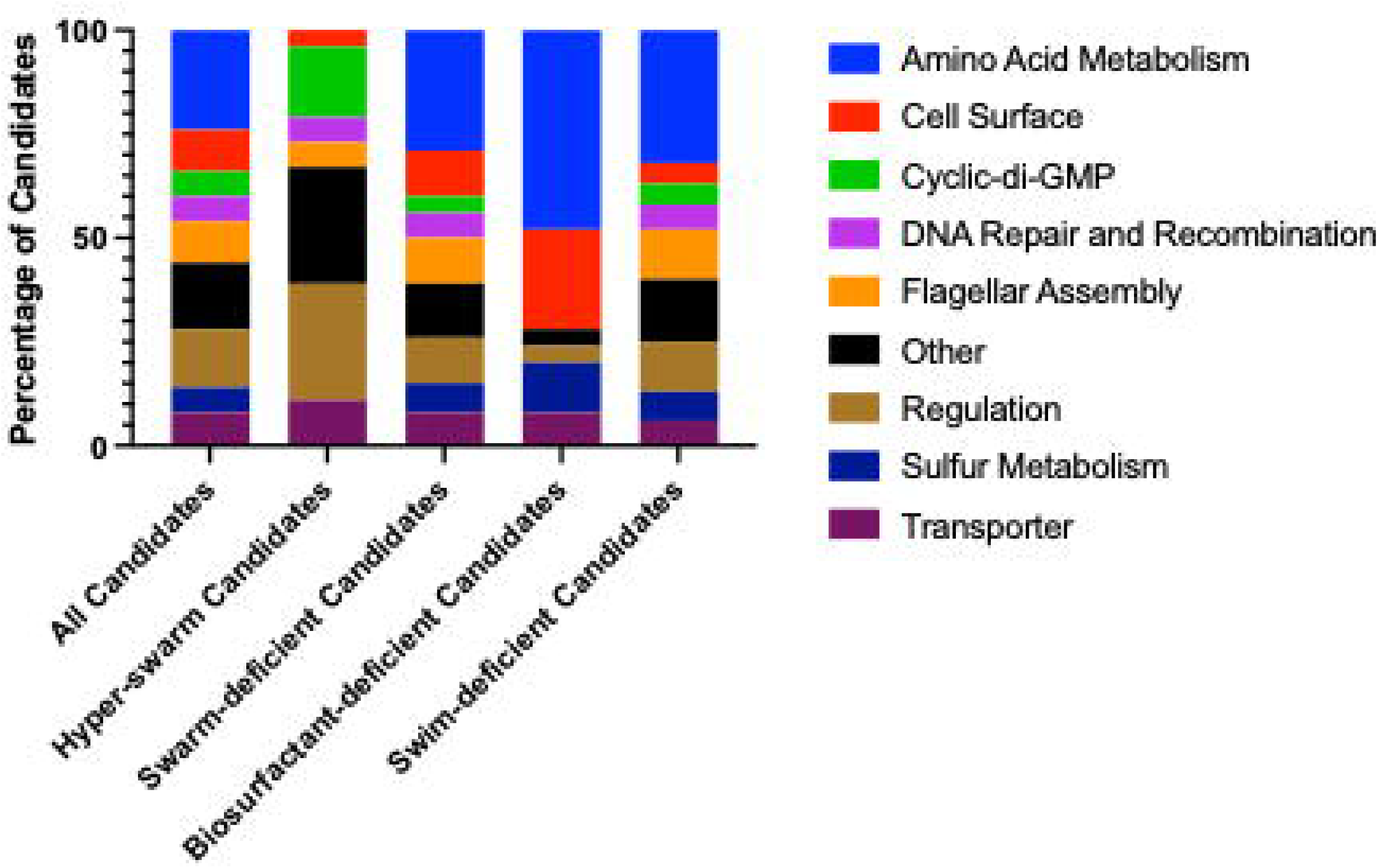
Summary of genetic loci identified by transposon mutagenesis. Stacked bar charts categorizing the candidates (n=108) identified from the transposon mutagenesis. Candidates were phenotypically sub-grouped based on swarming motility into hyper-swarming (n= 18) or swarm-deficient (n=90). The swarm-deficient group was then sub-grouped based on a loss of biosurfactant production (n=25) or a defect in swimming motility (n=82), with a subset of candidates present in both groups.

### Loss of genes related to Gacamide A biosynthesis and secretion result in loss of swarming motility and changes in biosurfactant production

We identified a class of mutants defective for both swarming motility and biosurfactant production (Table S4), but swim proficient. These mutants all had transposon insertions that mapped to a biosynthetic operon reported to encode the machinery needed to produce the non-ribosomal cyclic lipopeptide biosurfactant Gacamide A (34, 43). Gacamide A can promote swarming motility when purified and exogenously supplemented to the swarm-deficient parental *P. fluorescens* Pf0-1 (43, 44). The *gamA*, *gamB*, and *gamC* genes encode the biosynthetic proteins for Gacamide A. The *pleA*, *pleB*, and *pleC* genes are predicted to encode the Gacamide A secretion system given their sequence similarity to the MacAB-TolC type macrolide efflux pumps and conservation amongst cyclic lipopeptide-producing pseudomonads (43, 45–47).

Given these identified candidate genes, we conducted a genetic analysis utilizing the swarm-proficient Δ*rsmA* Δ*rsmE* Δ*rsmI* strain and introduced additional single chromosomal deletions of the predicted biosynthetic (*gamA*, *gamB*, and *gamC*) genes and transport (*pleA*, *pleB*, and *pleC*) genes. For these operonic genes, the individual deletion constructs were designed such that the relevant promoters remained intact to drive the expression of any downstream genes. These strains were then probed for biosurfactant production and swarm motility.

For the genes predicted to encode functions required for Gacamide A production, deletion of *gamA* resulted in a marked decrease in biosurfactant production (Figure 4A) and swarming motility (Figure 4B), deletion of *gamB* resulted in a modest, but statistically significant decrease in biosurfactant production (Figure 4C) and swarming motility (Figure 4D), and deletion of *gamC* resulted in a complete loss of biosurfactant production (Figure 4E) and swarming motility (Figure 4F). For the genes predicted to encode functions required for Gacamide A export, loss of the predicted periplasmic or outer membrane components encoded by *pleA* and *pleC* gene, respectively, resulted in the complete loss of the biosurfactant zone (Figure 5A and 5E, respectively) and swarming motility (Figure 5B and 5F, respectively). Interestingly, loss of the predicted inner membrane component encoded by *pleB* did not result in a reduction of biosurfactant zone (Figure 5C) but did result in a sizeable decrease in swarming motility (Figure 5D).

**FIG 4.**
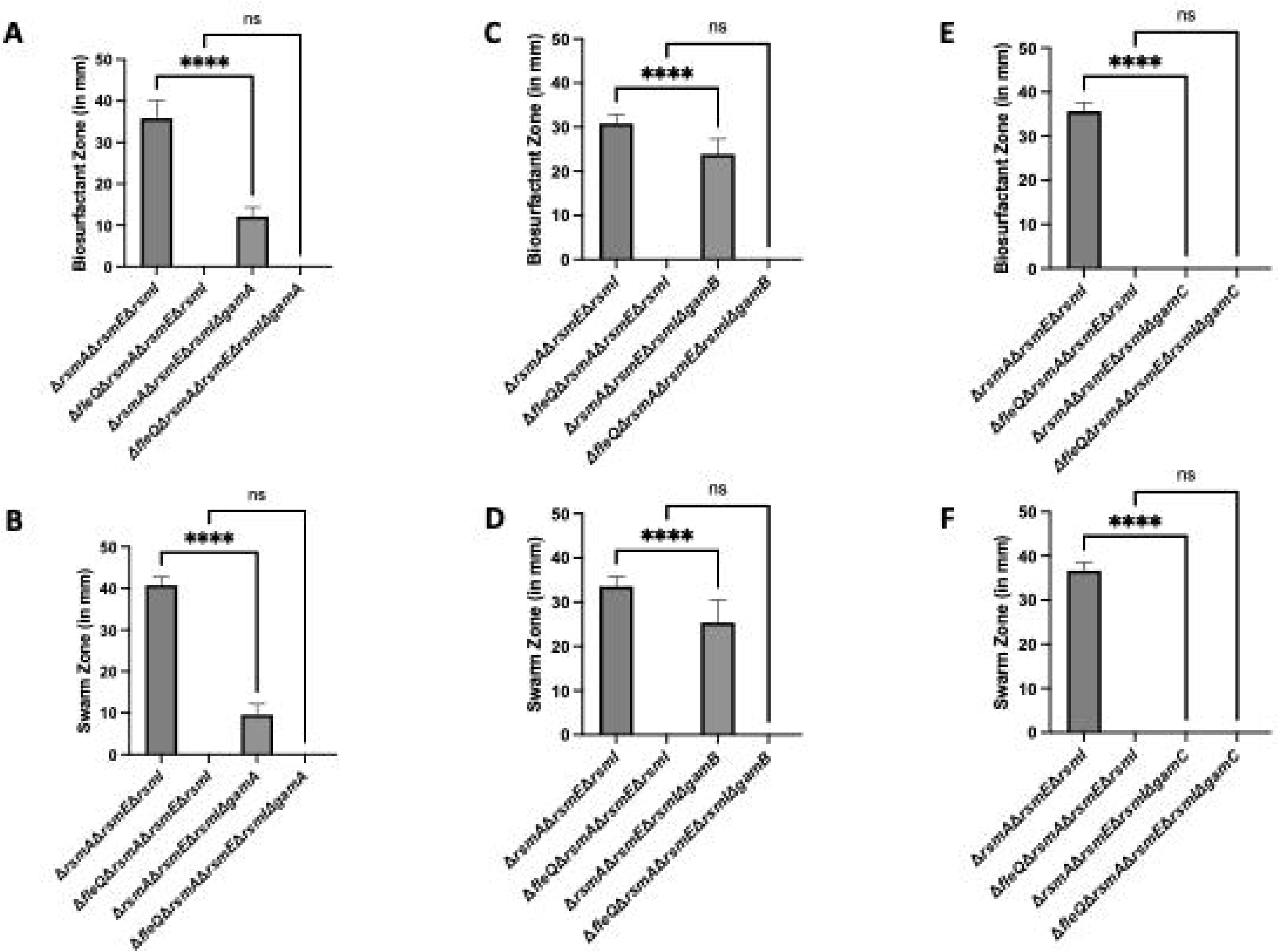
Loss of components of the biosurfactant operon effect swarming motility through the altered biosurfactant levels. (A) Biosurfactant zone (in millimeters) of the Δ*rsmA* Δ*rsmE* Δ*rsmI* triple mutant, Δ*fleQ* Δ*rsmA* Δ*rsmE* Δ*rsmI* quadruple mutant, Δ*rsmA* Δ*rsmE* Δ*rsmI* Δ*gamA* quadruple mutant, and the Δ*fleQ* Δ*rsmA* Δ*rsmE* Δ*rsmI* Δ*gamA* quintuple mutant after inoculation of 2.5µl of overnight culture on the surface of KA minimal medium supplemented with 0.5% agar followed by 24h growth at 30°C. (B) Swarm zone (in millimeters) of the Δ*rsmA* Δ*rsmE* Δ*rsmI* triple mutant, Δ*fleQ* Δ*rsmA* Δ*rsmE* Δ*rsmI* quadruple mutant, Δ*rsmA* Δ*rsmE* Δ*rsmI* Δ*gamA* quadruple mutant, and the Δ*fleQ* Δ*rsmA* Δ*rsmE* Δ*rsmI* Δ*gamA* quintuple mutant after inoculation of 2.5µl of overnight culture on the surface of KA minimal medium supplemented with 0.5% agar followed by 24h growth at 30°C then 24h growth at room temperature. (C) Biosurfactant zone (in millimeters) of the Δ*rsmA* Δ*rsmE* Δ*rsmI* triple mutant, Δ*fleQ* Δ*rsmA* Δ*rsmE* Δ*rsmI* quadruple mutant, Δ*rsmA* Δ*rsmE* Δ*rsmI* Δ*gamB* quadruple mutant, and the Δ*fleQ* Δ*rsmA* Δ*rsmE* Δ*rsmI* Δ*gamB* quintuple mutant after inoculation of 2.5µl of overnight culture on the surface of KA minimal medium supplemented with 0.5% agar followed by 24h growth at 30°C. (D) Swarm zone (in millimeters) of the Δ*rsmA* Δ*rsmE* Δ*rsmI* triple mutant, Δ*fleQ* Δ*rsmA* Δ*rsmE* Δ*rsmI* quadruple mutant, Δ*rsmA* Δ*rsmE* Δ*rsmI* Δ*gamB* quadruple mutant, and the Δ*fleQ* Δ*rsmA* Δ*rsmE* Δ*rsmI* Δ*gamB* quintuple mutant after inoculation of 2.5µl of overnight culture on the surface of KA minimal medium supplemented with 0.5% agar followed by 24h growth at 30°C then 24h growth at room temperature . (E) Biosurfactant zone (in millimeters) of the Δ*rsmA* Δ*rsmE* Δ*rsmI* triple mutant, Δ*fleQ* Δ*rsmA* Δ*rsmE* Δ*rsmI* quadruple mutant, Δ*rsmA* Δ*rsmE* Δ*rsmI* Δ*gamC* quadruple mutant, and the Δ*fleQ* Δ*rsmA* Δ*rsmE* Δ*rsmI* Δ*gamC* quintuple mutant after inoculation of 2.5µl of overnight culture on the surface of KA minimal medium supplemented with 0.5% agar followed by 24h growth at 30°C. (F) Swarm zone (in millimeters) of the Δ*rsmA* Δ*rsmE* Δ*rsmI* triple mutant, Δ*fleQ* Δ*rsmA* Δ*rsmE* Δ*rsmI* quadruple mutant, Δ*rsmA* Δ*rsmE* Δ*rsmI* Δ*gamC* quadruple mutant, and the Δ*fleQ* Δ*rsmA* Δ*rsmE* Δ*rsmI* Δ*gamC* quintuple mutant after inoculation of 2.5µl of overnight culture on the surface of KA minimal medium supplemented with 0.5% agar followed by 24h growth at 30°C then 24h growth at room temperature. Statistical significance for this figure was determined using one-way ANOVA with Tukey’s multiple comparisons tests. ****, P<0.0001. All error bars represent standard deviation. All error bars represent standard deviation.

**FIG 5.**
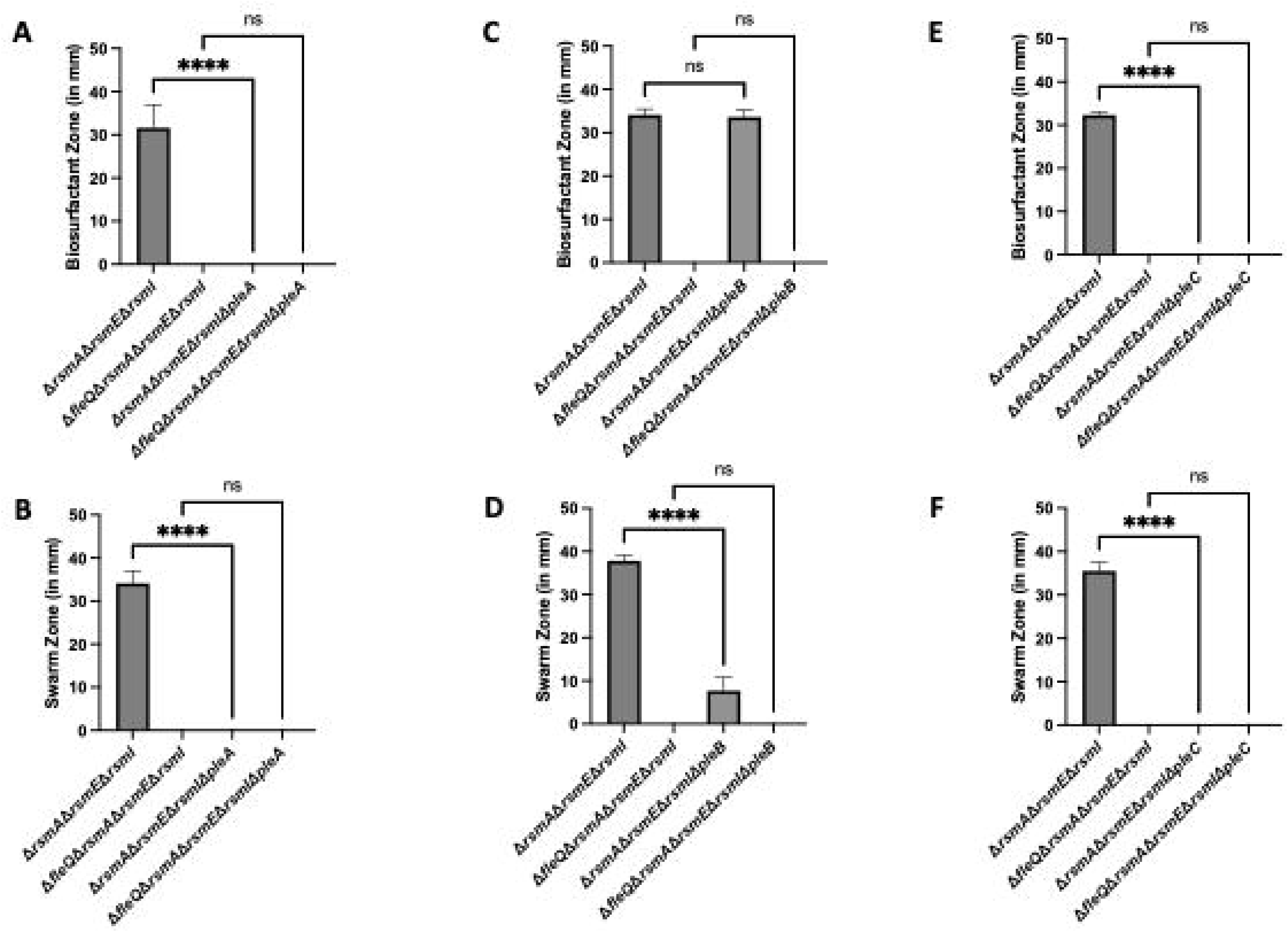
Loss of any component of the biosurfactant secretion system results in a swarm motility defect. (A) Biosurfactant zone (in millimeters) of the Δ*rsmA* Δ*rsmE* Δ*rsmI* triple mutant, Δ*fleQ* Δ*rsmA* Δ*rsmE* Δ*rsmI* quadruple mutant, Δ*rsmA* Δ*rsmE* Δ*rsmI* Δ*pleA* quadruple mutant, and the Δ*fleQ* Δ*rsmA* Δ*rsmE* Δ*rsmI* Δ*pleA* quintuple mutant after inoculation of 2.5µl of overnight culture on the surface of KA minimal medium supplemented with 0.5% agar followed by 24h growth at 30°C. (B) Swarm zone (in millimeters) of the Δ*rsmA* Δ*rsmE* Δ*rsmI* triple mutant, Δ*fleQ* Δ*rsmA* Δ*rsmE* Δ*rsmI* quadruple mutant, Δ*rsmA* Δ*rsmE* Δ*rsmI* Δ*pleA* quadruple mutant, and the Δ*fleQ* Δ*rsmA* Δ*rsmE* Δ*rsmI* Δ*pleA* quintuple mutant after inoculation of 2.5µl of overnight culture on the surface of KA minimal medium supplemented with 0.5% followed by 24h growth at 30°C then 24h growth at room temperature. (C) Biosurfactant zone (in millimeters) of the Δ*rsmA* Δ*rsmE* Δ*rsmI* triple mutant, Δ*fleQ* Δ*rsmA* Δ*rsmE* Δ*rsmI* quadruple mutant, Δ*rsmA* Δ*rsmE* Δ*rsmI* Δ*pleB* quadruple mutant, and the Δ*fleQ* Δ*rsmA* Δ*rsmE* Δ*rsmI* Δ*pleB* quintuple mutant after inoculation of 2.5µl of overnight culture on the surface of KA minimal medium supplemented with 0.5% agar followed by 24h growth at 30°C. (D) Swarm zone (in millimeters) of the Δ*rsmA* Δ*rsmE* Δ*rsmI* triple mutant, Δ*fleQ* Δ*rsmA* Δ*rsmE* Δ*rsmI* quadruple mutant, Δ*rsmA* Δ*rsmE* Δ*rsmI* Δ*pleB* quadruple mutant, and the Δ*fleQ* Δ*rsmA* Δ*rsmE* Δ*rsmI* Δ*pleB* quintuple mutant after inoculation of 2.5µl of overnight culture on the surface of KA minimal medium supplemented with 0.5% agar followed by 24h growth at 30°C then 24h growth at room temperature. (E) Biosurfactant zone (in millimeters) of the Δ*rsmA* Δ*rsmE* Δ*rsmI* triple mutant, Δ*fleQ* Δ*rsmA* Δ*rsmE* Δ*rsmI* quadruple mutant, Δ*rsmA* Δ*rsmE* Δ*rsmI* Δ*pleC* quadruple mutant, and the Δ*fleQ* Δ*rsmA* Δ*rsmE* Δ*rsmI* Δ*pleC* quintuple mutant after inoculation of 2.5µl of overnight culture on the surface of KA minimal medium supplemented with 0.5% agar followed by 24h growth at 30°C. (F) Swarm zone (in millimeters) of the Δ*rsmA* Δ*rsmE* Δ*rsmI* triple mutant, Δ*fleQ* Δ*rsmA* Δ*rsmE* Δ*rsmI* quadruple mutant, Δ*rsmA* Δ*rsmE* Δ*rsmI* Δ*pleC* quadruple mutant, and the Δ*fleQ* Δ*rsmA* Δ*rsmE* Δ*rsmI* Δ*pleC* quintuple mutant after inoculation of 2.5µl of overnight culture on the surface of KA minimal medium supplemented with 0.5% agar followed by 24h growth at 30°C then 24h growth at room temperature. Statistical significance was determined using one-way ANOVA with Tukey’s multiple comparisons tests. ****, P<0.0001. All error bars represent standard deviation.

To confirm that the observed differences in swarming motility were not related to changes in flagellar function, we assessed the strains for their ability to swim. After 24h of growth at 30°C on swim agar, all tested mutants produced swim zones similar to their parental strain (Figure S5 and S6). The similar swim zones also indicated that there was no general growth defect for any of these mutants.

### Loss of FliA results in a swarming and swimming motility defect

We identified several candidate mutants defective for both swimming and swarming motility. We focused our analysis specifically on the alternative sigma factor FliA, which was previously associated with flagellar biosynthesis and swarming motility regulation in *P. aeruginosa* (48, 49). We made a chromosomal deletion of *fliA* in the swarm-proficient Δ*rsmA* Δ*rsmE* Δ*rsmI* strain and assessed the mutant for the ability to swim and swarm.

After 24h of growth at 30°C on swim agar, the Δ*rsmA* Δ*rsmE* Δ*rsmI* Δ*fliA* strain was completely deficient in swimming motility (Figure 6A). After 24h of growth at 30°C and 24h of growth at room temperature on swarm agar, the Δ*rsmA* Δ*rsmE* Δ*rsmI* Δ*fliA* strain was completely deficient in swarming motility (Figure 6B). We noted no significant difference in biosurfactant production between the Δ*rsmA* Δ*rsmE* Δ*rsmI* Δ*fliA* mutant and its parental strain (Figure S7), indicating that loss of FliA function results in loss of motility via loss of flagellar function.

**FIG 6.**
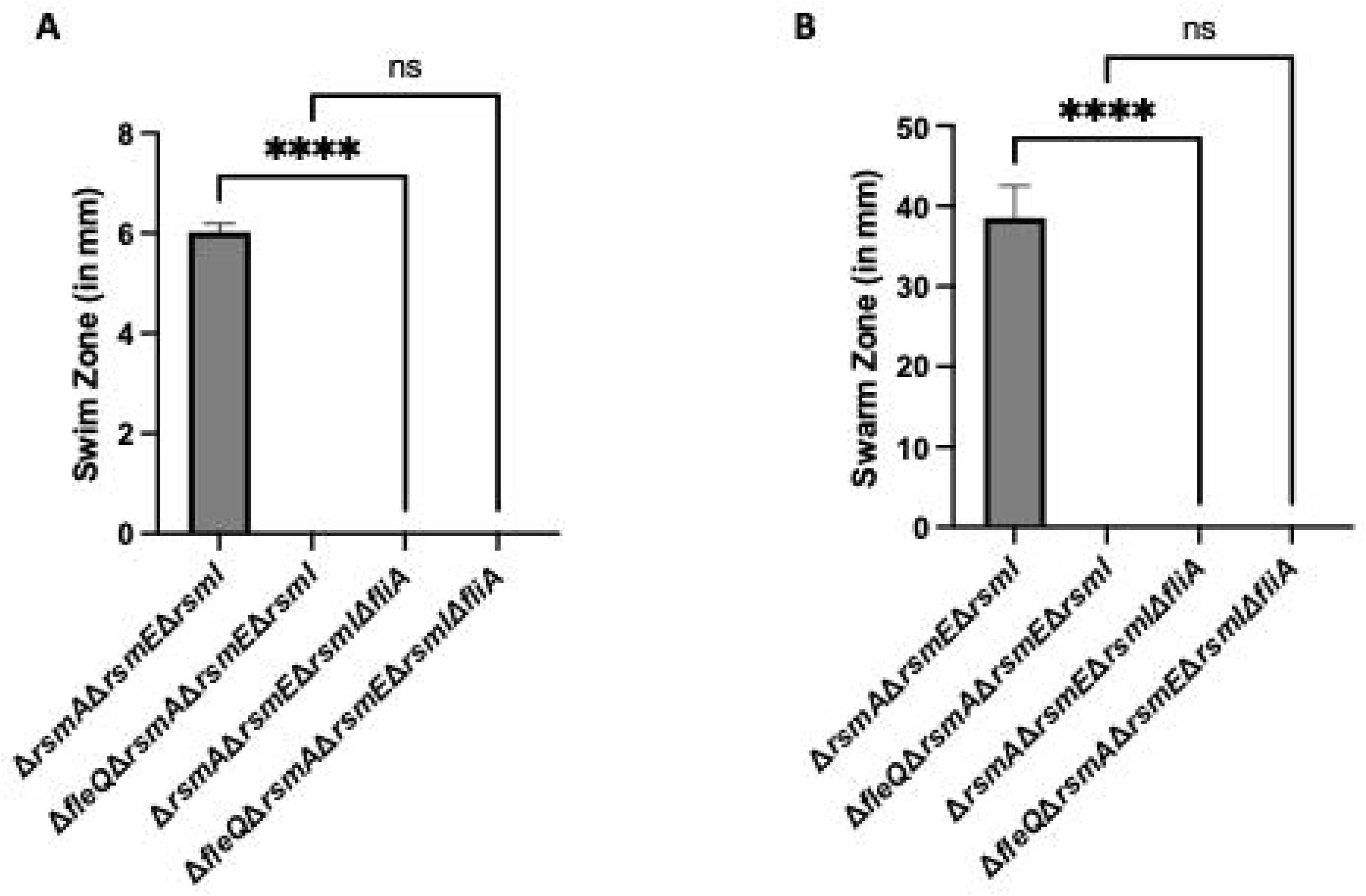
A FliA-deficient strain has a motility defect likely due to loss of flagellar function. (A) Swim zone (in millimeters) of the Δ*rsmA* Δ*rsmE* Δ*rsmI* triple mutant, Δ*fleQ* Δ*rsmA* Δ*rsmE* Δ*rsmI* quadruple mutant, Δ*rsmA* Δ*rsmE* Δ*rsmI* Δ*fliA* quadruple mutant, and the Δ*fleQ* Δ*rsmA* Δ*rsmE* Δ*rsmI* Δ*fliA* quintuple mutant after toothpick inoculation on KA minimal medium supplemented with 0.3% agar followed by 24h growth at 30°C. (B) Swarm zone (in millimeters) of the Δ*rsmA* Δ*rsmE* Δ*rsmI* triple mutant, Δ*fleQ* Δ*rsmA* Δ*rsmE* Δ*rsmI* quadruple mutant, Δ*rsmA* Δ*rsmE* Δ*rsmI* Δ*fliA* quadruple mutant, and the Δ*fleQ* Δ*rsmA* Δ*rsmE* Δ*rsmI* Δ*fliA* quintuple mutant after inoculation of 2.5µl of overnight culture on the surface of KA minimal medium supplemented with 0.5% agar followed by 24h growth at 30°C then 24h growth at room temperature. Statistical significance for this figure was determined using one-way ANOVAs with Tukey’s multiple comparisons tests. ****, P<0.0001. All error bars represent standard deviation.

## Discussion

Our data here build upon the link between the Gac/Rsm pathway and swarming motility reported by our group and several other teams (22, 32, 34, 35, 40, 43, 44). Our genetic analyses demonstrate that loss of the Rsm proteins, which was previously shown to phenotypically mimic overstimulation of the Gac/Rsm pathway (36), induces biosurfactant production/secretion and swarming motility of *P. fluorescens* Pf0-1 and does so in a RsmA- and RsmE-dependent manner. These findings support previously published data showing that a *P. fluorescens* Pf0-1 merodiploid strain containing the native Pf0-1 *gacA* gene and non-native *gacA* gene from *P. protogens* Pf-5 was proficient for swarming motility (34) and build on previous results demonstrating that RsmE regulates biosurfactant secretion by *P. fluorescens* Pf0-1 (44).

Our subsequent transposon mutagenesis identified numerous genes that contribute to swarming motility via the production of a functional flagellum, biosurfactant biosynthesis/ secretion, or both. The mutants with a defect in biosurfactant production included strains with transposon insertions in genes encoding the non-ribosomal peptide synthetases responsible for synthesizing the biosurfactant Gacamide A and its predicted outer membrane transporter PleC. Subsequent genetic analyses revealed that loss of *gamA*, *gamB*, or *gamC* decreased biosurfactant secretion and swarming motility to varying degrees. These differences could be attributed to the likelihood that loss of the *gamA* or *gamB* gene still results in the production of basal levels of Gacamide A or an alternative surfactant that can partially support swarming motility. Evidence supporting this second idea has been previously reported by other groups, whereby domain deletions and substitutions within non-ribosomal peptide synthetases results in an alternative peptide being synthesized, which is a strategy currently being utilized to develop novel biotherapeutics (50–55). Based on the observed complete loss of swarming motility but the ability of this strain to swim, a mutation in the *gamC* gene likely results in complete loss of biosurfactant production. The predicted terminal tandem thioesterase (TE-TE) domain of GamC is likely required for circularization and release of the synthesized peptide from its peptidyl carrier protein (43, 56, 57). Interestingly, previously published mutations in *gamA* via antibiotic resistance cassette (34) or mini-Tn5 insertion (44) resulted in complete loss of biosurfactant secretion whereas a chromosomal deletion resulted in a sizeable decrease but not complete loss of biosurfactant secretion. We posit that disruption of the *gamA* gene via insertional mutagenesis has the unintended consequence of inactivating the whole operon, whereas chromosomal deletion likely allows for the continued production of the GamB and GamC proteins.

Genetic analysis revealed that the predicted Gacamide A secretion machinery is also required for swarming motility. As expected, loss of the periplasmic component PleA or the outer membrane component PleC of the proposed Gacamide A secretion machinery resulted in loss of detectable biosurfactant and complete loss of swarming motility. Interestingly, loss of the inner-membrane component PleB results in complete loss of swarming motility even though this mutant strain is apparently proficient for biosurfactant production and swimming motility. These observations could suggest the presence of an alternative inner membrane transporter component that can substitute for PleB to support the secretion of Gacamide A, or that PleB may modify Gacamide A during secretion, which may bne required for this surfactant to promote swarming motility.

The identified candidates with defects only in swimming and swarming motility clustered to a variety of seemingly unrelated pathways. Since multiple studies will be required to fully investigate the role of these many functions in motility, we focused here specifically on the alternative sigma factor FliA. As expected based on other studies in pseudomonads (48, 49, 58), a loss of FliA resulted in the loss of swimming and swarming motility, likely via loss of flagellar function.

Collectively, our findings provide evidence that swarming motility is regulated via the Gac/Rsm pathway in an RsmA- and RsmE-dependent manner by impacting flagellar motility and biosurfactant production/secretion. These findings provide a set of tools for the future study of these pathways, and we hope the findings from the transposon screen may be used by the broader community to study both motility and other Gac/Rsm-dependent phenotypes of *P. fluorescens* Pf0-1.

## Materials and Methods

### Strains and media used in this study

*P. fluorescens* Pf0-1, *E. coli* S17-1 λ-pir, and *E. coli* SM10 λ-pir were used throughout this study. *E. coli* and *P. fluorescens* were routinely grown in lysogeny broth (LB) and *P. fluorescens* was routinely grown in KA minimal medium, as previously defined by Collins et al. (59). KA minimal medium contains 50 mM Tris-HCl (pH 7.4), 0.61 mM MgSO_4_, 1 mM K_2_HPO_4_, and 0.4% (wt/vol) L-arginine HCl. Medium was supplemented with 30µg/ml gentamycin for *P. fluorescens* when harboring the expression vector pMQ72. For *E. coli*, medium was supplemented with 10 ug/ml gentamycin when harboring the allelic exchange vector pMQ30 or the expression vector pMQ72, with 50 µg/ml carbenicillin for the strain harboring the transposon-containing shuttle vector pBT20, or 15 ug/ml tetracycline when harboring the allelic exchange plasmid pEX18Tc. The strains and plasmids used in this study are listed in **Table S6**.

### Construction of in-frame chromosomal gene deletions

For chromosomal deletions in this study, the allelic exchange vectors pEX18Tc or pMQ30 were utilized. Flanking regions of the target genes were amplified via PCR using Phusion polymerase (New England BioLabs). All primers used in the study are listed in **Table S7**.

The amplicons were integrated into SmaI (New England BioLabs)-digested vector using the GeneArt Gibson Assembly HiFi Master Mix (Invitrogen) according to the manufacturer specifications. Constructs were electroporated into *E. coli* S17-1 λ-pir and plated on LB agar with 15 µg/ml tetracycline or 10 µg/ml gentamycin for pEX18Tc or pMQ30-based constructs respectively. Integration of the flanking regions into the vector was confirmed via PCR amplification and Sanger sequencing. Constructs were conjugated into *P. fluorescens*, whereby 1mL aliquots of *E. coli* harboring the construct and *P. fluorescens* cultures grown overnight at 30°C were mixed in a 2 mL microcentrifuge tube, pelleted, washed in LB, and then plated on LB agar with no antibiotic selection to facilitate uptake of the deletion construct by *P. fluorescens*. After overnight incubation at 30°C, the cells were scraped from the plate and resuspended in fresh LB liquid medium. Serial dilutions were plated on LB agar supplemented with 45 µg/ml tetracycline and 30 µg/ml chloramphenicol for pEX18TC-based constructs or 30 µg/ml gentamycin and 30 µg/ml chloramphenicol for pMQ30-based constructs to select and counter select, respectively, for integration of the constructs into the *P. fluorescens* genome. To facilitate looping out of the drug resistance cassette, merodiploid candidates were initially grown in LB liquid without antibiotic selection overnight at 30°C and then serial dilutions were plated on LB agar without sodium chloride supplemented with 10% sucrose. Single colonies were struck on LB agar with and without antibiotic selection to confirm loss of the resistance cassette carried by the plasmid. Antibiotic susceptible candidates were screened for loss of the target gene via PCR amplification and Sanger sequencing.

### Construction of the *rsmA* expression vector

The *rsmA* gene was PCR amplified using Phusion polymerase (New England BioLabs) according to the manufacturer specifications. The primers were designed such that the forward primer contained the high-affinity T7 phage gene 10 ribosomal binding site (60) and 8bp spacer nucleotides upstream of the *rsmA* start codon. Additionally, both primers contained 20bp of homology to the pMQ72 SmaI cut-site at their 5’ ends. Primers used to build this construct are listed in **Table S7**.

The amplicon was integrated into SmaI-digested pMQ72 using the GeneArt Gibson Assembly HiFi Master Mix (Invitrogen) according to the manufacturer specifications. The construct was electroporated into *E. coli* S17-1 λ-pir, cells were recovered in LB for 1h at 30°C, and subsequently plated on LB agar supplemented with 10 µg/ml gentamycin. Antibiotic-resistant candidates were sequenced to confirm gene insertion into pMQ72. The construct was then electroporated into *P. fluorescens*, cells were initially recovered in LB for 1h at 30°C, and dilutions were plated on LB agar supplemented with 30 µg/ml gentamycin to select for retention of the construct.

### Swim assay

Swim assays were conducted as previously defined in Pastora and O’Toole (36). Briefly, *P. fluorescens* Pf0-1 strains were grown overnight in 5 ml of LB with appropriate antibiotic selection at 30°C hours with agitation. 1mL aliquots were transferred to sterile 1.5 mL microcentrifuge tubes and used to stab inoculate KA minimal medium supplemented with 0.3% agar (swim agar). Inoculated plates were incubated for 24 hours at 30°C. The diameter of resulting swim zones was measured using a ruler.

### Biosurfactant and swarm assays

Biosurfactant and swarm assays were conducted using a modified approach previously described by Ha et al. (61). Briefly, *P. fluorescens* Pf0-1 strains were grown overnight in 5 ml of LB with appropriate antibiotic selection at 30°C hours with agitation. 2.5µl of each culture was pipetted directly on the surface of KA minimal medium supplemented with 0.5% agar (swarm agar). Plates were initially incubated for 24 hours at 30°C and the diameter of the resulting biosurfactant zones was measured using a ruler. Plates were then incubated for an additional 24 hours at room temperature and the diameter of the resulting swarm zones was measured using a ruler.

### Transposon mutagenesis

Transposon mutagenesis was conducted as previously defined in Pastora and O’Toole (36). Briefly, *E. coli* harboring the pBT20 plasmid and the *P. fluorescens* Pf0-1 *ΔrsmAΔrsmEΔrsmI* triple mutant were grown in LB, with appropriate antibiotic selection, overnight at 30°C and mixed in a 1:1 ratio in a 2mL microcentrifuge tube. Cell pellets were then washed, resuspended, and plated on LB agar to facilitate conjugation of the pBT20 plasmid into *P. fluorescens*. Plates were incubated for 90 min at 30°C, and the resulting cells were scraped from the plate, washed in fresh LB, serially diluted, and plated on LB agar supplemented with 30 µg/ml gentamycin and 30 ug/ml chloramphenicol to select for transposon integration in *P. fluorescens*. Per Pastora and O’Toole (36), mutant libraries were generated by picking single colonies with sterile pipette tips into sterile 96-well flat-bottom polystyrene plates. Candidates were subsequently screened for their ability to swarm on swarm agar (see above). Candidates that were unable to swarm or were able to swarm at the 24h timepoint and continue to swarm at the 48h timepoint (hyper-swarmers) were identified from the mutant library. The identified candidates were struck out from the mutant library onto LB agar supplemented with 30 µg/ml gentamycin and subsequently re-tested for swarming motility to verify the swarm-deficient or hyper-swarm phenotype. Verified candidates were subsequently phenotypically screened for swimming motility and biosurfactant secretion. Transposon chromosomal insertion sites and transposon orientations were determined using arbitrary primed PCR as described by O’Toole et al. (62).

## Data availability

This manuscript has no large datasets.

## Acknowledgement

We would like to thank Dr. Fabrice Jean-Pierre and Dr. Sherry Kuchma for many helpful discussions about swarming motility by *Pseudomonas aeruginosa*. Sanger sequencing was provided by the Dartmouth Molecular Biology Core. This work was supported by NIH grant R01AI168017 to G.A.O. The Dartmouth Molecular Biology Core was supported by NIH grant P30CA023108 to the Dartmouth Cancer Center.

