## Supplemental Figures for "Multiple Pathways Impact Swarming Motility of *Pseudomonas fluorescens* Pf0-1"

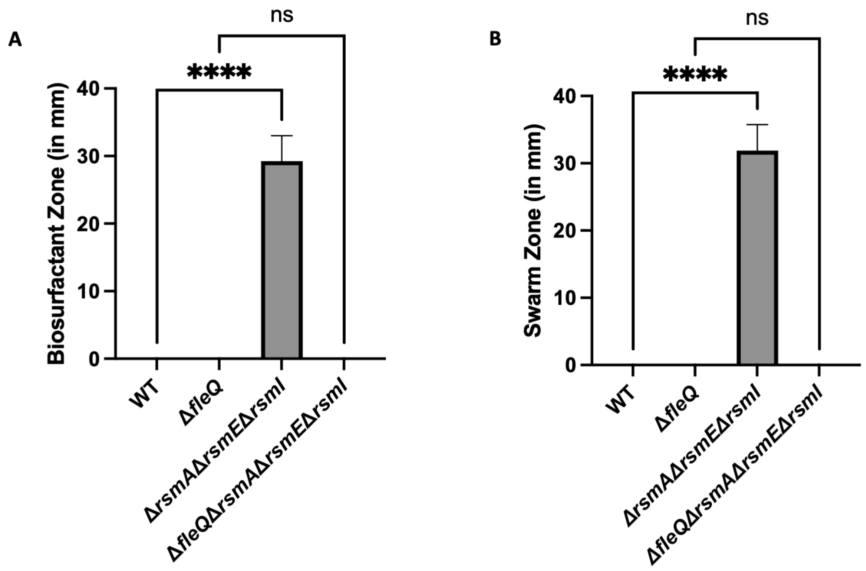


**FIG S1 A strain deficient for the Rsm proteins swarms on 0.3% Agar.** (A) Biosurfactant zone (in millimeters) of the WT strain, Δ*fleQ* single mutant, Δ*rsmA* Δ*rsmE* Δ*rsmI* triple mutant, and a Δ*fleQ* Δ*rsmA* Δ*rsmE* Δ*rsmI* quadruple mutant after inoculation of 2.5µl of overnight culture on the surface of KA minimal medium supplemented with 0.3% agar (swim agar) and incubated for 24h at 30°C. (B) Swarm zone (in millimeters) of the WT strain, Δ*fleQ* single mutant, Δ*rsmA* Δ*rsmE* Δ*rsmI* triple mutant, and a Δ*fleQ* Δ*rsmA* Δ*rsmE* Δ*rsmI* quadruple mutant after inoculation of 2.5µl of overnight culture on the surface of KA minimal medium supplemented with 0.3% agar (swim agar) followed by 24h growth at 30°C followed by 24h growth at room temperature. Statistical significance for this figure was determined using one-way ANOVA with Tukey's multiple comparisons tests. ****, P<0.0001. All error bars represent standard deviation.

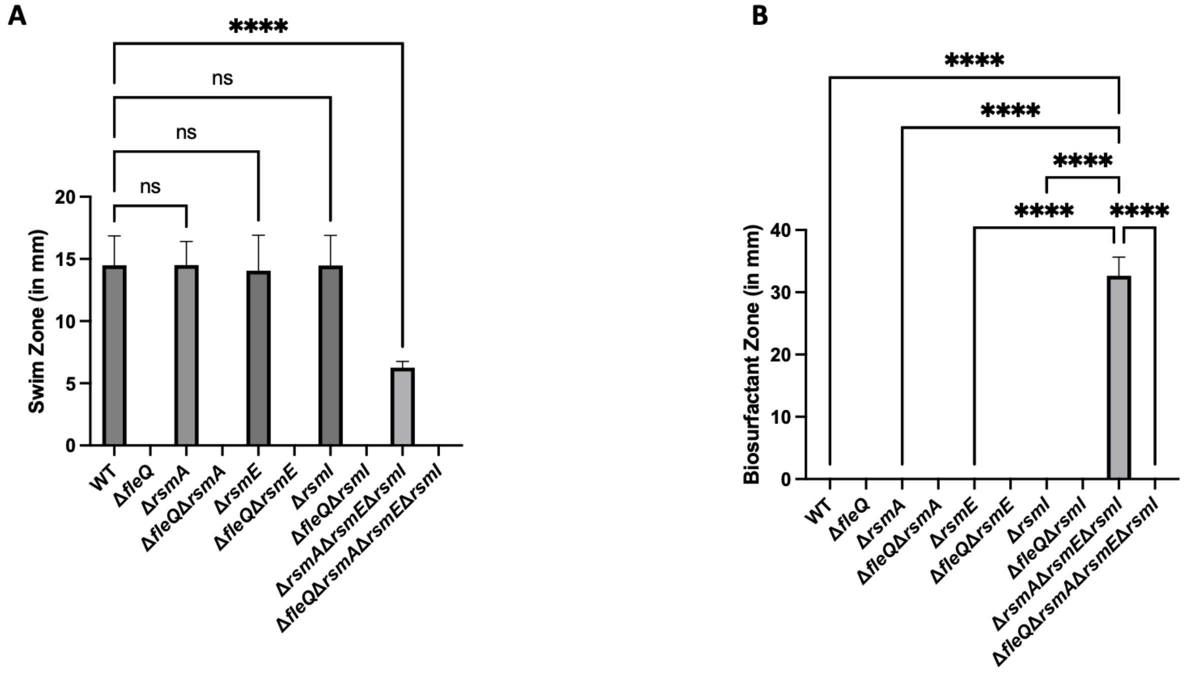


**FIG S2 Loss of a single Rsm Protein is not sufficient for biosurfactant production**. (A) Swim zone (in millimeters) of the WT strain, Δ*fleQ* single mutant, Δ*rsmA* single mutant, Δ*fleQ* Δ*rsmA* double mutant, Δ*rsmE* single mutant, Δ*fleQ* Δ*rsmE* double mutant, Δ*rsmI* single mutant, Δ*fleQ* Δ*rsmI* double mutant, Δ*rsmA* Δ*rsmE* Δ*rsmI* triple mutant, and a Δ*fleQ* Δ*rsmA* Δ*rsmE* Δ*rsmI* quadruple mutant after toothpick inoculation on KA minimal medium supplemented with 0.3% agar and 24h growth at 30°C. (B) Biosurfactant zone (in millimeters) of the WT strain, Δ*fleQ* single mutant, Δ*rsmA* single mutant, Δ*fleQ* Δ*rsmA* double mutant, Δ*rsmE* single mutant, Δ*fleQ* Δ*rsmE* double mutant, Δ*rsmI* single mutant, Δ*fleQ* Δ*rsmI* double mutant, Δ*rsmA* Δ*rsmE* Δ*rsmI* triple mutant, and a Δ*fleQ* Δ*rsmA* Δ*rsmE* Δ*rsmI* quadruple mutant after inoculation of 2.5µl of overnight culture on the surface of KA minimal medium supplemented with 0.5% agar and 24h growth at 30°C. Statistical significance for this figure was determined using one-way ANOVA with Tukey's multiple comparisons tests. ****, $P<0.0001$. All error bars represent standard deviation.

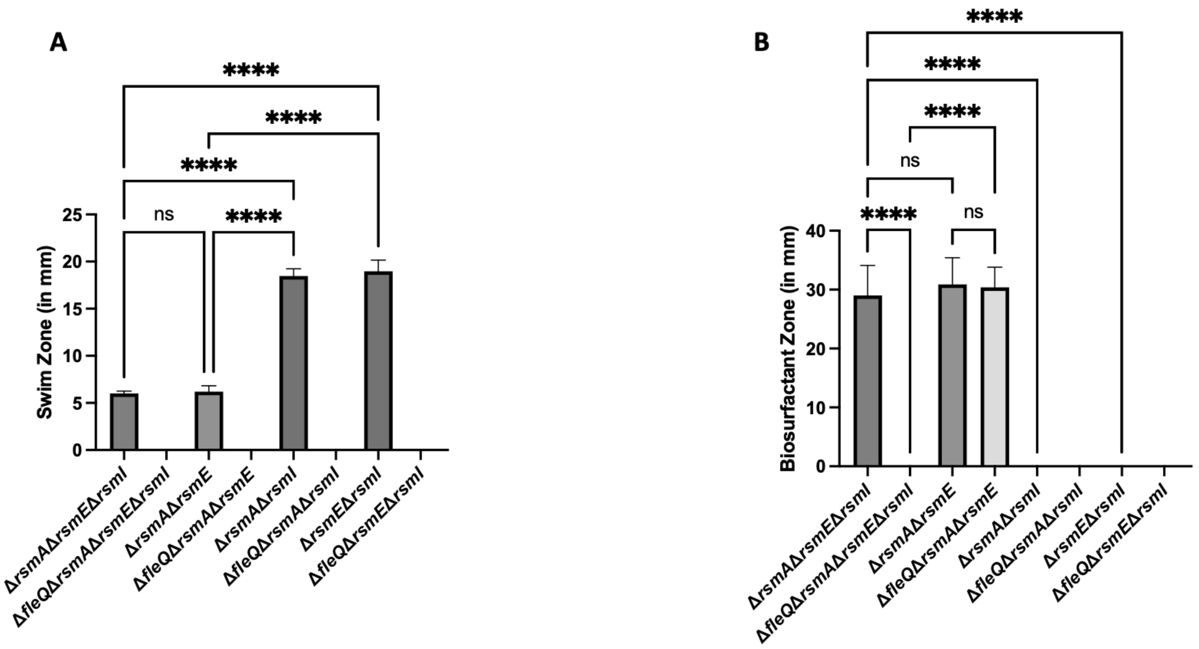


**FIG S3 A strain deficient for both RsmA and RsmE is sufficient for biosurfactant production.** (A) Swim zone (in millimeters) of the Δ*rsmA* Δ*rsmE* Δ*rsmI* triple mutant, Δ*fleQ* Δ*rsmA* Δ*rsmE* Δ*rsmI* quadruple mutant, Δ*rsmA* Δ*rsmE* double mutant, Δ*fleQ* Δ*rsmA* Δ*rsmE* triple mutant, Δ*rsmA* Δ*rsmI* double mutant, Δ*fleQ* Δ*rsmA* Δ*rsmI* triple mutant, Δ*rsmE* Δ*rsmI* double mutant, and Δ*fleQ* Δ*rsmE* Δ*rsmI* triple mutant after toothpick inoculation on KA minimal medium supplemented with 0.3% agar and 24h growth at 30°C. (B) Biosurfactant zone (in millimeters) the Δ*rsmA* Δ*rsmE* Δ*rsmI* triple mutant, Δ*fleQ* Δ*rsmA* Δ*rsmE* Δ*rsmI* quadruple mutant, Δ*rsmA* Δ*rsmE* double mutant, Δ*fleQ* Δ*rsmA* Δ*rsmE* triple mutant, Δ*rsmA* Δ*rsmI* double mutant, Δ*fleQ* Δ*rsmA* Δ*rsmI* triple mutant, Δ*rsmE* Δ*rsmI* double mutant, and Δ*fleQ* Δ*rsmE* Δ*rsmI* triple mutant after inoculation of 2.5µl of overnight culture on the surface of KA minimal medium supplemented with 0.5% agar and 24h growth at 30°C. Statistical significance for this figure was determined using one-way ANOVA with Tukey's multiple comparisons tests. ****, $P<0.0001$. All error bars represent standard deviation.

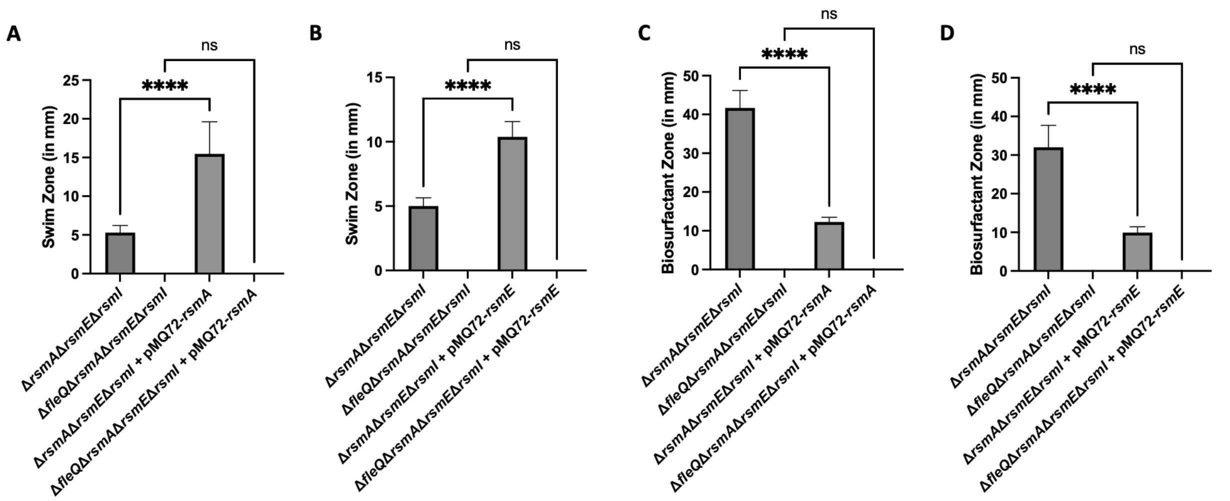


**FIG S4 Expression of RsmA or RsmE from a plasmid reduces biosurfactant production in a strain deficient of the Rsm proteins.** (A) Swim zone (in millimeters) of the Δ*rsmA* Δ*rsmE* Δ*rsmI* mutant, Δ*fleQ* Δ*rsmA* Δ*rsmE* Δ*rsmI* mutant, Δ*rsmA* Δ*rsmE* Δ*rsmI* mutant + pMQ72-*rsmA*, and Δ*fleQ* Δ*rsmA* Δ*rsmE* Δ*rsmI* + pMQ72-*rsmA* mutant after toothpick inoculation on KA minimal medium supplemented with 0.3% agar and 24h growth at 30°C. (B) Swim zone (in millimeters) of the Δ*rsmA* Δ*rsmE* Δ*rsmI* mutant, Δ*fleQ* Δ*rsmA* Δ*rsmE* Δ*rsmI* mutant, Δ*rsmA* Δ*rsmE* Δ*rsmI* mutant + pMQ72-*rsmE*, and Δ*fleQ* Δ*rsmA* Δ*rsmE* Δ*rsmI* + pMQ72-*rsmE* mutant after toothpick inoculation on KA minimal medium supplemented with 0.3% agar and 24h growth at 30°C. (C) Biosurfactant zone (in millimeters) of the Δ*rsmA* Δ*rsmE* Δ*rsmI* mutant, Δ*fleQ* Δ*rsmA* Δ*rsmE* Δ*rsmI* mutant, Δ*rsmA* Δ*rsmE* Δ*rsmI* mutant + pMQ72-*rsmA*, and Δ*fleQ* Δ*rsmA* Δ*rsmE* Δ*rsmI* + pMQ72-*rsmA* mutant after inoculation of 2.5µl of overnight culture on the surface of KA minimal medium supplemented with 0.5% agar and 24h growth at 30°C. (D) Biosurfactant zone (in millimeters) of the Δ*rsmA* Δ*rsmE* Δ*rsmI* mutant, Δ*fleQ* Δ*rsmA* Δ*rsmE* Δ*rsmI* mutant, Δ*rsmA* Δ*rsmE* Δ*rsmI* mutant + pMQ72-*rsmE*, and Δ*fleQ* Δ*rsmA* Δ*rsmE* Δ*rsmI* + pMQ72-*rsmE* mutant after inoculation of 2.5µl of overnight culture on the surface of KA minimal medium supplemented with 0.5% agar and 24h growth at 30°C. Statistical significance for this figure was determined using one-way ANOVA with Tukey's multiple comparisons tests. ****, P<0.0001. All error bars represent standard deviation.

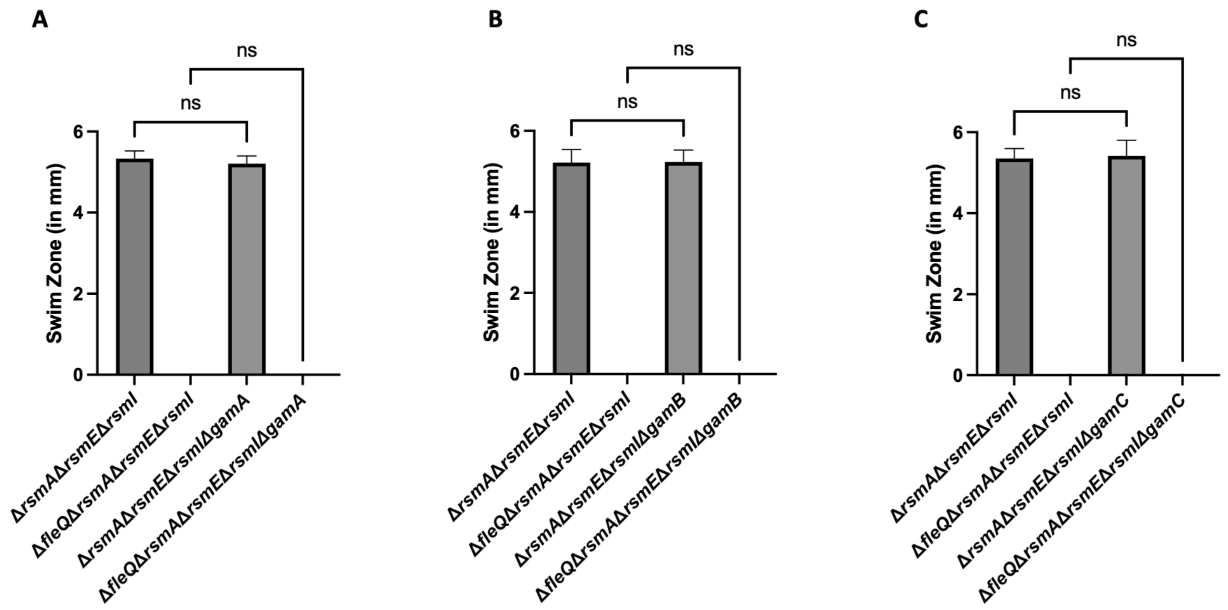


**FIG S5 Loss of any component of the predicted biosurfactant biosynthetic machinery does not affect flagellar function as assessed by swimming motility.** (A) Swim zone (in millimeters) of the Δ*rsmA* Δ*rsmE* Δ*rsmI* triple mutant, Δ*fleQ* Δ*rsmA* Δ*rsmE* Δ*rsmI* quadruple mutant, Δ*rsmA* Δ*rsmE* Δ*rsmI* Δ*gamA* quadruple mutant, and the Δ*fleQ* Δ*rsmA* Δ*rsmE* Δ*rsmI* Δ*gamA* quintuple mutant after toothpick inoculation on KA minimal medium supplemented with 0.3% agar and 24h growth at 30°C. (B) Swim zone (in millimeters) of the Δ*rsmA* Δ*rsmE* Δ*rsmI* triple mutant, Δ*fleQ* Δ*rsmA* Δ*rsmE* Δ*rsmI* quadruple mutant, Δ*rsmA* Δ*rsmE* Δ*rsmI* Δ*gamB* quadruple mutant, and the Δ*fleQ* Δ*rsmA* Δ*rsmE* Δ*rsmI* Δ*gamB* quintuple mutant after toothpick inoculation on KA minimal medium supplemented with 0.3% agar and 24h growth at 30°C. (C) Swim zone (in millimeters) of the Δ*rsmA* Δ*rsmE* Δ*rsmI* triple mutant, Δ*fleQ* Δ*rsmA* Δ*rsmE* Δ*rsmI* quadruple mutant, Δ*rsmA* Δ*rsmE* Δ*rsmI* Δ*gamC* quadruple mutant, and the Δ*fleQ* Δ*rsmA* Δ*rsmE* Δ*rsmI* Δ*gamC* quintuple mutant after toothpick inoculation on KA minimal medium supplemented with 0.3% agar and 24h growth at 30°C. Statistical significance for this figure was determined using one-way ANOVA with Tukey's multiple comparisons tests. All error bars represent standard deviation.

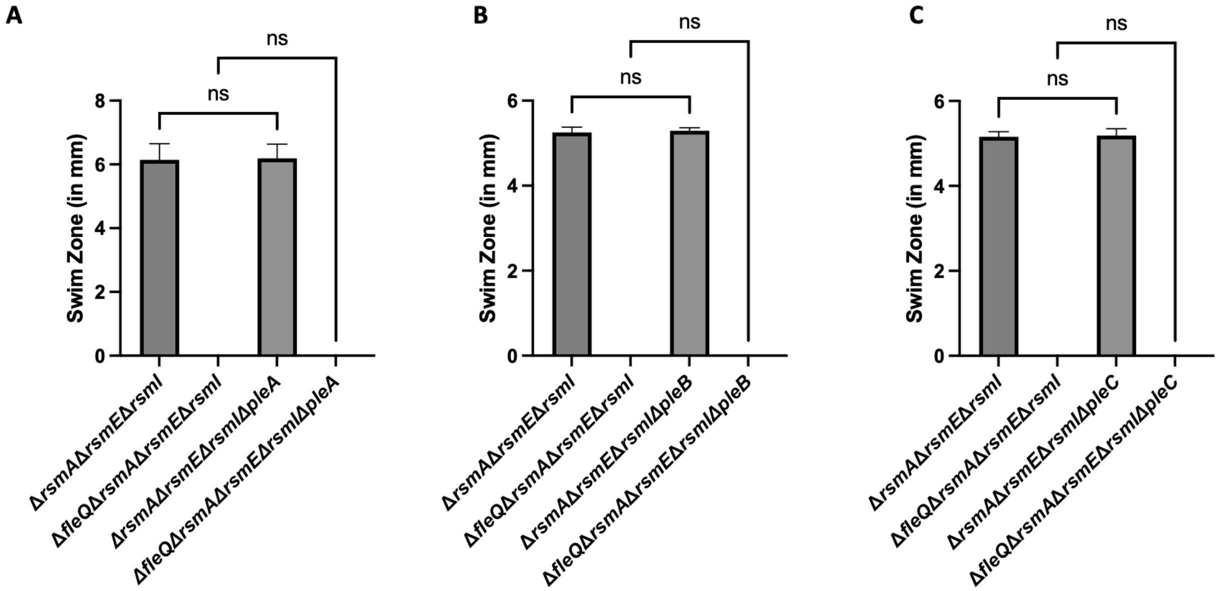


**FIG S6 Loss of any component of the predicted biosurfactant secretion machinery does not affect flagellar function as assessed by swimming motility.** (A) Swim zone (in millimeters) of the Δ*rsmA* Δ*rsmE* Δ*rsmI* triple mutant, Δ*fleQ* Δ*rsmA* Δ*rsmE* Δ*rsmI* quadruple mutant, Δ*rsmA* Δ*rsmE* Δ*rsmI* Δ*pleA* quadruple mutant, and the Δ*fleQ* Δ*rsmA* Δ*rsmE* Δ*rsmI* Δ*pleA* quintuple mutant after toothpick inoculation on KA minimal medium supplemented with 0.3% agar and 24h growth at 30°C. (B) Swim zone (in millimeters) of the Δ*rsmA* Δ*rsmE* Δ*rsmI* triple mutant, Δ*fleQ* Δ*rsmA* Δ*rsmE* Δ*rsmI* quadruple mutant, Δ*rsmA* Δ*rsmE* Δ*rsmI* Δ*pleB* quadruple mutant, and the Δ*fleQ* Δ*rsmA* Δ*rsmE* Δ*rsmI* Δ*pleB* quintuple mutant after toothpick inoculation on KA minimal medium supplemented with 0.3% agar and 24h growth at 30°C. (C) Swim zone (in millimeters) of the Δ*rsmA* Δ*rsmE* Δ*rsmI* triple mutant, Δ*fleQ* Δ*rsmA* Δ*rsmE* Δ*rsmI* quadruple mutant, Δ*rsmA* Δ*rsmE* Δ*rsmI* Δ*pleC* quadruple mutant, and the Δ*fleQ* Δ*rsmA* Δ*rsmE* Δ*rsmI* Δ*pleC* quintuple mutant after toothpick inoculation on KA minimal medium supplemented with 0.3% agar and 24h growth at 30°C. Statistical significance for this figure was determined using one-way ANOVA with Tukey's multiple comparisons tests. All error bars represent standard deviation.

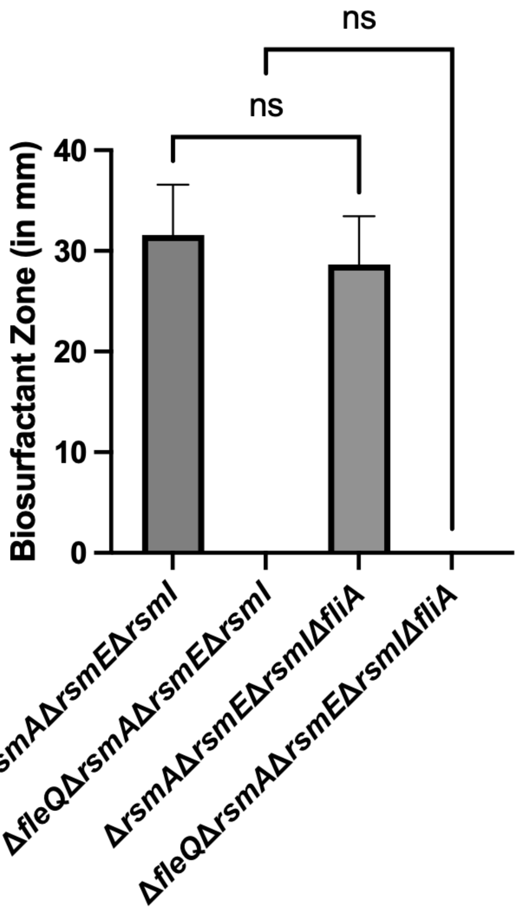


**FIG S7 Loss of FliA has no impact on biosurfactant production.** Biosurfactant zone (in millimeters) of the Δ*rsmA* Δ*rsmE* Δ*rsmI* triple mutant, Δ*fleQ* Δ*rsmA* Δ*rsmE* Δ*rsmI* quadruple mutant, Δ*rsmA* Δ*rsmE* Δ*rsmI* Δ*fliA* quadruple mutant, and the Δ*fleQ* Δ*rsmA* Δ*rsmE* Δ*rsmI* Δ*fliA* quintuple mutant after inoculation of 2.5µl of overnight culture on the surface of KA minimal medium supplemented with 0.5% agar and 24h growth at 30°C. Statistical significance was determined using a one-way ANOVA with Tukey's multiple comparisons tests. All error bars represent standard deviation.
