## Supplemental Tables S6-7 for "Multiple Pathways Impact Swarming Motility of *Pseudomonas fluorescens* Pf0-1"

**Table S6. Strains and plasmids used in this study.**

| Strain/Plasmid | Genotype/Description | Reference/Source |
|---|---|---|
| ***Escherichia coli*** | | |
| S17-1 (λpir) | recA, thi, pro, hsdR- M+RP4::2-Tc::Mu::Km Tn7 $Tp^R$ $Sm^R$ λpir | (1) |
| SM10 (λpir) | thi, thr, leu, tonA, lacY, supE, recA::RP4-2-$Tc^R$::Mu Km λpir | (2) |
| ***Pseudomonas fluorescens* Pf0-1** | | |
| SMC 4798 | LapA-HA; HA tag inserted after residue 4093 | (3) |
| SMC 8581 | MapA-HA; HA tag inserted after residue 2601 | (4) |
| SMC 9499 | Δ*fleQ* LapA-HA | (5) |
| SMC 8746 | Δ*fleQ* MapA-HA | (5) |
| SMC 9502 | Δ*rsmA* LapA-HA | (5) |
| SMC 9503 | Δ*rsmA*Δ*fleQ* LapA-HA | (5) |
| SMC 9507 | Δ*rsmE* LapA-HA | (5) |
| SMC 9508 | Δ*rsmE*Δ*fleQ* LapA-HA | (5) |
| SMC 9610 | Δ*rsmI* LapA-HA | (5) |
| SMC 9611 | Δ*rsmI*Δ*fleQ* LapA-HA | (5) |
| SMC 9858 | Δ*rsmA*Δ*rsmE* LapA-HA | This Study |
| SMC 9859 | Δ*rsmA*Δ*rsmE*Δ*fleQ* LapA-HA | This Study |
| SMC 9614 | Δ*rsmA*Δ*rsmI* LapA-HA | (5) |
| SMC 9615 | Δ*rsmA*Δ*rsmI*Δ*fleQ* LapA-HA | (5) |
| SMC 9860 | Δ*rsmE*Δ*rsmI* LapA-HA | This study |

| | | |
|---|---|---|
| SMC 9861 | Δ*rsmE*Δ*rsmI*Δ*fleQ* LapA-HA | This study |
| SMC 9612 | Δ*rsmA*Δ*rsmE*Δ*rsmI* LapA-HA | (5) |
| SMC 9613 | Δ*rsmA*Δ*rsmE*Δ*rsmI*Δ*fleQ* LapA-HA | (5) |
| SMC 9862 | Δ*rsmA*Δ*rsmE*Δ*rsmI*Δ*fliA* LapA-HA | This study |
| SMC 9863 | Δ*rsmA*Δ*rsmE*Δ*rsmI*Δ*fleQ*Δ*fliA* LapA-HA | This study |
| SMC 9864 | Δ*rsmA*Δ*rsmE*Δ*rsmI*Δ*gamA* LapA-HA | This study |
| SMC 9865 | Δ*rsmA*Δ*rsmE*Δ*rsmI*Δ*fleQ*Δ*gamA* LapA-HA | This study |
| SMC 9866 | Δ*rsmA*Δ*rsmE*Δ*rsmI*Δ*gamB* LapA-HA | This study |
| SMC 9867 | Δ*rsmA*Δ*rsmE*Δ*rsmI*Δ*fleQ*Δ*gamB* LapA-HA | This study |
| SMC 9868 | Δ*rsmA*Δ*rsmE*Δ*rsmI*Δ*gamC* LapA-HA | This study |
| SMC 9869 | Δ*rsmA*Δ*rsmE*Δ*rsmI*Δ*fleQ*Δ*gamC* LapA-HA | This study |
| SMC 9870 | Δ*rsmA*Δ*rsmE*Δ*rsmI*Δ*pleA* LapA-HA | This study |
| SMC 9871 | Δ*rsmA*Δ*rsmE*Δ*rsmI*Δ*fleQ*Δ*pleA* LapA-HA | This study |
| SMC 9872 | Δ*rsmA*Δ*rsmE*Δ*rsmI*Δ*pleB* LapA-HA | This study |
| SMC 9873 | Δ*rsmA*Δ*rsmE*Δ*rsmI*Δ*fleQ*Δ*pleB* LapA-HA | This study |
| SMC 9874 | Δ*rsmA*Δ*rsmE*Δ*rsmI*Δ*pleC* LapA-HA | This study |
| SMC 9875 | Δ*rsmA*Δ*rsmE*Δ*rsmI*Δ*fleQ*Δ*pleC* LapA-HA | This study |
| SMC 9876 | Δ*rsmA*Δ*rsmE*Δ*rsmI*; pMQ72-*rsmA* | This study |
| SMC 9877 | Δ*rsmA*Δ*rsmE*Δ*rsmI*Δ*fleQ* LapA-HA; pMQ72-*rsmA* | This study |
| SMC 9702 | Δ*rsmA*Δ*rsmE*Δ*rsmI* LapA-HA; pMQ72-*rsmE* | (5) |
| SMC 9703 | Δ*rsmA*Δ*rsmE*Δ*rsmI*Δ*fleQ* LapA-HA; pMQ72-*rsmE* | (5) |

| **Plasmids** | | |
|---|---|---|
| SMC 1210 | pBT20 | (6) |
| SMC 560 | pEX18Tc | (7) |
| SMC 2765 | pMQ30 | (8) |
| SMC 2795 | pMQ72 | (8) |
| SMC 220 | pSMC21 | (9) |
| SMC 9850 | pEX18Tc Δ*gamA* | This study |
| SMC 9851 | pEX18Tc Δ*gamB* | This study |
| SMC 9852 | pEX18Tc Δ*gamC* | This study |
| SMC 9854 | pEX18Tc Δ*pleB* | This study |
| SMC 9855 | pEX18Tc Δ*pleC* | This study |
| SMC 7975 | pMQ30 Δ*fleQ* | (5) |
| SMC 9849 | pMQ30 Δ*fliA* | This study |
| SMC 9853 | pMQ30 Δ*pleA* | This study |
| SMC 9492 | pMQ30 Δ*rsmA* | (10) |
| SMC 9506 | pMQ30 Δ*rsmE* | (5) |
| SMC 9642 | pMQ30 Δ*rsmI* | (5) |
| SMC 9856 | pMQ72-*rsmA* | This study |
| SMC 9699 | pMQ72-*rsmE* | (5) |

**Table S7. Primers used in this study.**

| Primer Name | Primer Sequence (5'-3') | Purpose |
|---|---|---|
| del fliA F1 | gtcgactctagaggatccccgtgagcatggacagtttccgtatcg | Amplify region upstream of *fliA* |
| del fliA R1 | cccacactgccttcatttcatagcacaggtcctgccg | Amplify region upstream of *fliA* |
| del fliA F2 | aggacctgtgctatgaaatgaaggcagtgtggggacac | Amplify region downstream of *fliA* |
| del fliA R2 | cgaattcgagctcggtacccgcgaaaaattgcgttgagcagatctg | Amplify region downstream of *fliA* |
| fliA seq F | caatctgttcgccaagttgaccaag | Verify Deletion of *fliA* |
| fliA seq R | gttggtgaacccgagatcacg | Verify Deletion of *fliA* |
| del gamA F1 | gtcgactctagaggatccccgtgcgcgtggcgctcgat | Amplify region upstream of *gamA* |
| del gamA R1 | gtcggctcagagcaagatccatctcacgtgatttggcgc | Amplify region upstream of *gamA* |
| del gamA F2 | aatcacgtgagatggatcttgctctgagccgacaccgg | Amplify region downstream of *gamA* |
| del gamA R2 | cgaattcgagctcggtaccccatcagttgcgcgtcttcgaagtc | Amplify region downstream of *gamA* |
| gamA seq F | tcgtgactgtacggtctagtg | Verify Deletion of *gamA* |
| gamA seq R | gggtggaaataacagccatcag | Verify Deletion of *gamA* |
| del gamB F1 | gtcgactctagaggatcccctgctggacgcccacgatc | Amplify region upstream of *gamB* |
| del gamB R1 | cagttcgatcacgttcacgggggtaacctgctgtaaattcgg | Amplify region upstream of *gamB* |
| del gamB F2 | cagcaggttacccccgtgaacgtgatcgaactgttggc | Amplify region downstream of *gamB* |

| | | |
|---|---|---|
| del gamB R2 | cgaattcgagctcggtacccggccaggtggaacagactgg | Amplify region downstream of *gamB* |
| gamB seq F | gaccaggcaaaacagctggaag | Verify Deletion of *gamB* |
| gamB seq R | gtcacctgatcaagttcggcc | Verify Deletion of *gamB* |
| del gamC F1 | gtcgactctagaggatcccccctcgctcatggtgacacc | Amplify region upstream of *gamC* |
| del gamC R1 | gataaaaacttcaatcaggtcacaggacgatctccattctttcc | Amplify region upstream of *gamC* |
| del gamC F2 | gagatcgtcctgtgacctgattgaagtttttatcctggaaccc | Amplify region downstream of *gamC* |
| del gamC R2 | cgaattcgagctcggtaccccggaatcgtcagcaccgatttg | Amplify region downstream of *gamC* |
| gamC seq F | gatcagagtgctgccatcacc | Verify Deletion of *gamC* |
| gamC seq R | ccagccactgacccttcttg | Verify Deletion of *gamC* |
| del pleA F1 | gtcgactctagaggatccccctacgaagtcgcgatgcaattgctc | Amplify region upstream of *pleA* |
| del pleA R1 | aggttcgctcatgcaaatgaccacaattccgttcagggttg | Amplify region upstream of *pleA* |
| del pleA F2 | cggaattgtggtcatttgcatgagcgaacctctgctg | Amplify region downstream of *pleA* |
| del pleA R2 | cgaattcgagctcggtaccccagcatcagtttgttgacgttgtcttc | Amplify region downstream of *pleA* |
| pleA seq F | cgacgctggacgcatcg | Verify Deletion of *pleA* |
| pleA seq R | gcgctgaaagataaagccgaaatagtc | Verify Deletion of *pleA* |
| del pleB F1 | gtcgactctagaggatccccgacgagagcacgcaaatgcg | Amplify region upstream of *pleB* |
| del pleB R1 | cgccggtcagtcgctcatgcccctgccactttc | Amplify region upstream of *pleB* |
| del pleB F2 | caggggcatgagcgactgaccggcgggcatg | Amplify region downstream of *pleB* |

| | | |
|---|---|---|
| del pleB R2 | cgaattcgagctcggtaccccaaaatcacgccatgagaccggttaatatg | Amplify region downstream of *pleB* |
| pleB seq F | caaatcggtgctgacgattccg | Verify Deletion of *pleB* |
| pleB seq R | gaaccgtgagatggaagtgtgc | Verify Deletion of *pleB* |
| del pleC F1 | gtcgactctagaggatcccccagttcgacatagccaccgatg | Amplify region upstream of *pleC* |
| del pleC R1 | aaggtgtaagaaaaaatgaccaaacagcctcgttcacgaatgag | Amplify region upstream of *pleC* |
| del pleC F2 | gaacgaggctgtttggtcattttttcttacaccttttgtcgccag | Amplify region downstream of *pleC* |
| del pleC R2 | cgaattcgagctcggtacccggcactaattcgggagggtatctg | Amplify region downstream of *pleC* |
| pleC seq F | catcaccttgtcagtgcgtgag | Verify Deletion of *pleC* |
| pleC seq R | cttgccgtacttgtacgacagttc | Verify Deletion of *pleC* |
| pMQ72 *rsmA* F | cgaattcgagctcggtacccgaaggagatatacatatgctgattctgactcgtcggtg | Amplify *rsmA* |
| pMQ72 *rsmA R* | gtcgactctagaggatccccttaatggcttggttcttcgtccttc | Amplify *rsmA* |
| pMQ72 *rsmE* F | cgaattcgagctcggtacccgaaggagatatacatatgctgatactcacccgcaaagtc | Amplify *rsmE* |
| pMQ72 *rsmE* R | gtcgactctagaggatcccctcagggcgtttgtggcttgtc | Amplify *rsmE* |
| pMQ30 seq F | gatgcctggcagttccctactctcg | Sequencing pMQ30/pEX18Tc MCS |
| pMQ30 seq R | ctggcacgacaggtttcccgactg | Sequencing pMQ30/pEX18Tc MCS |
| pMQ72 seq F | gaattgtgagcggataacaatttcacacag | Sequencing pMQ72 MCS |
| pMQ72 seq R | gacaactccagtgaaaagttcttctcctttac | Sequencing pMQ72 MCS |
| Rnd1_pBT20Tn | tataatgtgtggaattgtgagcgg | Arbitrary PCR |
| Rnd2_pBT20Tn | acaggaaacaggactctagagg | Arbitrary PCR |
| Seq_pBT20Tn | cacccagctttcttgtacac | Arbitrary PCR |
| GAO_ARB1 | ggccacgcgtcgactagtacnnnnnnnnnngatat | Arbitrary PCR |

| | | |
|---|---|---|
| GAO_ARB2 | ggccacgcgtcgactagtac | Arbitrary PCR |
